# Assessing the basal gene expression of cancer cell lines for in vitro transcriptomic toxicology screening

**DOI:** 10.1101/2023.07.26.550714

**Authors:** Michael B. Black, Alina Y. Efremenko, Patrick D. McMullen, A. Rasim Barutcu, Andy Nong

## Abstract

In vitro toxicology has used immortalized cancer cell lines as model human systems for decades. However, these cell lines pose problems in designing toxicity testing programs as they inherently do not represent normal human biology. There is also a huge number of such cell lines to choose from, derived from human cancer cells from nearly every tissue. We explored the idea of using available basal gene expression data (NCI-60 cell line panel and Human GTEx tissue data) to determine if there was sufficient variability in cell line gene expression to group cell lines by relevance to specific human tissues. The transcriptomic analysis suggests that the variability in gene expression in cancer cell lines and in normal human tissue is minimal. The overall basal gene expression of cancer cells lines even overlapped normal human tissue gene expression. While some human tissues (e.g., lung) have basal expression profiles that do not appear to be like any cancer cell line, including cancers that may be derived from the same tissue, most human tissues show basal expression profiles comparable to several cancer cell lines. These results are important to address the genomic baseline and variability of cancer cell lines used for new approach methods of toxicity testing.

## INTRODUCTION

The isolation and cloning of HeLa cells from cervical cancer cells in 1951 ushered in a new era for in vitro biology cell models for studying cancer outside of the human body as well as devising medicines to combat cancers (Masters, 2002; Boehm and Hahn, 2004; Shoemaker, 2006; Lee et al., 2008; Stuelten et al., 2010; Savas et al., 2011; Niu and Wang, 2015; Beskow, 2016; Mirabelli et al., 2019). Today there are many immortalized cancer cell lines derived from cancers in nearly all human tissues. Thus, re-sampling of cells from a living source is unnecessary. Cancer cell lines have known culture requirements and are amenable to many forms of cell culture from simple 2D plating to more complex multi-cell and 3D model systems.

The National Cancer Institute developed its own panel of 60 cancer cell lines specifically for use in screening small molecules for anti-cancer properties (Chabner, 2016). Interest in the use of in vitro culture systems for toxicity testing was as far back as 1970’s, the same time frame as early developments in the use of immortalized cancer cell lines for medicine and drug development (Rees, 1980). While in vitro toxicology systems have seen continued development for decades now, there has been intense interest in cell-based systems in the past decade (Shukla et al., 2010; Singh et al., 2018). The increasing emphasis on non-animal testing, and the need to reduce toxicity screening costs, improving throughput, and developing more human-relevant biological systems has prompted an increasing interest in use of cell lines for in vitro toxicology. Immortalized cancer cell lines remain in wide-scale use for toxicology studies due to the wealth of information available for these in vitro models.

Whole cell transcriptomic studies are increasingly used for the advancement of new approach methods (NAM) in toxicology. As short-term, high throughput and low-cost methods, transcriptomics can quickly assess cellular mode of action (MOA) and compare responses among compounds. When combined with benchmark dose analyses, these studies can form the basis for relative potency analyses and deriving points of departure (POD) (Rowlands et al., 2013; Andersen et al., 2018; McMullen et al., 2019; Baltazar et al., 2020; Nault et al., 2020; Harrill et al., 2021). Combining toxicogenomic approaches and in vitro models offers one path forward to reduce or eliminate animal testing. Immortalized cancer cell lines are common models used for such studies.

An issue in designing in vitro transcriptomic studies is the selection of cell models. While immortalized cancer cell lines are often chosen for use in screening studies, the choice of specific cell lines is often problematic. Immortalized cancer cell lines are inherently not models of normal human biology as they are derived from cells experiencing abnormal biology (Roggen, 2011). A small, focused lead compound screening study may wish to target specific tissues depending on what is known, or not known, about a family of compounds. While a large screening program may be more concerned with covering as much of human tissue biology as possible to assess overall risk factors of many unrelated compounds.

For cell lines to be used in transcriptomic studies, we compared the expression profile of an unperturbed cell line to that of a healthy human tissue in an unperturbed state. Comparing these basal gene expression profiles may provide some insight into the suitability of a given cell line as a surrogate for a human tissue or tissues. The basis for this comparison already exists for normal human tissue in the data available through the Broad Institute’s Genotype-Tissue Expression (GTEx) project (Consortium, 2013; Carithers and Moore, 2015). Similarly, for immortalized cancer cell lines, the National Cancer Institutes has sequenced the 60 cell lines in its NCI-60 cell panel (Reinhold et al., 2019). These data were used to determine if there was sufficient difference in basal gene expression among the NCI-60 cell lines and degrees of correspondence with expression in the GTEx human tissues to sort cell lines and tissues into groups of similar basal expression profiles.

## RESULTS

A summary of the relationship between expression in the NCI-60 cancer cell lines and GTEx human tissues (Figure 1). This analysis shows two patterns of expression profiles between cancer cell lines and human tissue samples. The NCI-60 cell lines show a few individual cell lines with wide dispersion of individual cell lines (the leukemia cell lines RPMI-8226 and MOLT-4) as a group based on originating tissue. However, in the center mass of the NCI-60 PCA plot we see one or more cell lines from all nine tissues of origin clustered with overlapping 95% confidence ellipses. Regarding GTEx human tissues, while the 14 brain tissues show a large 95% confidence ellipse, they all remain distinct from the other tissues. There is a group in the center of Figure 1(B) with minimal separation. This included colon tissues, arterial tissues, esophagus, adipose, adrenal gland, and bladder.

**Figure 1.**
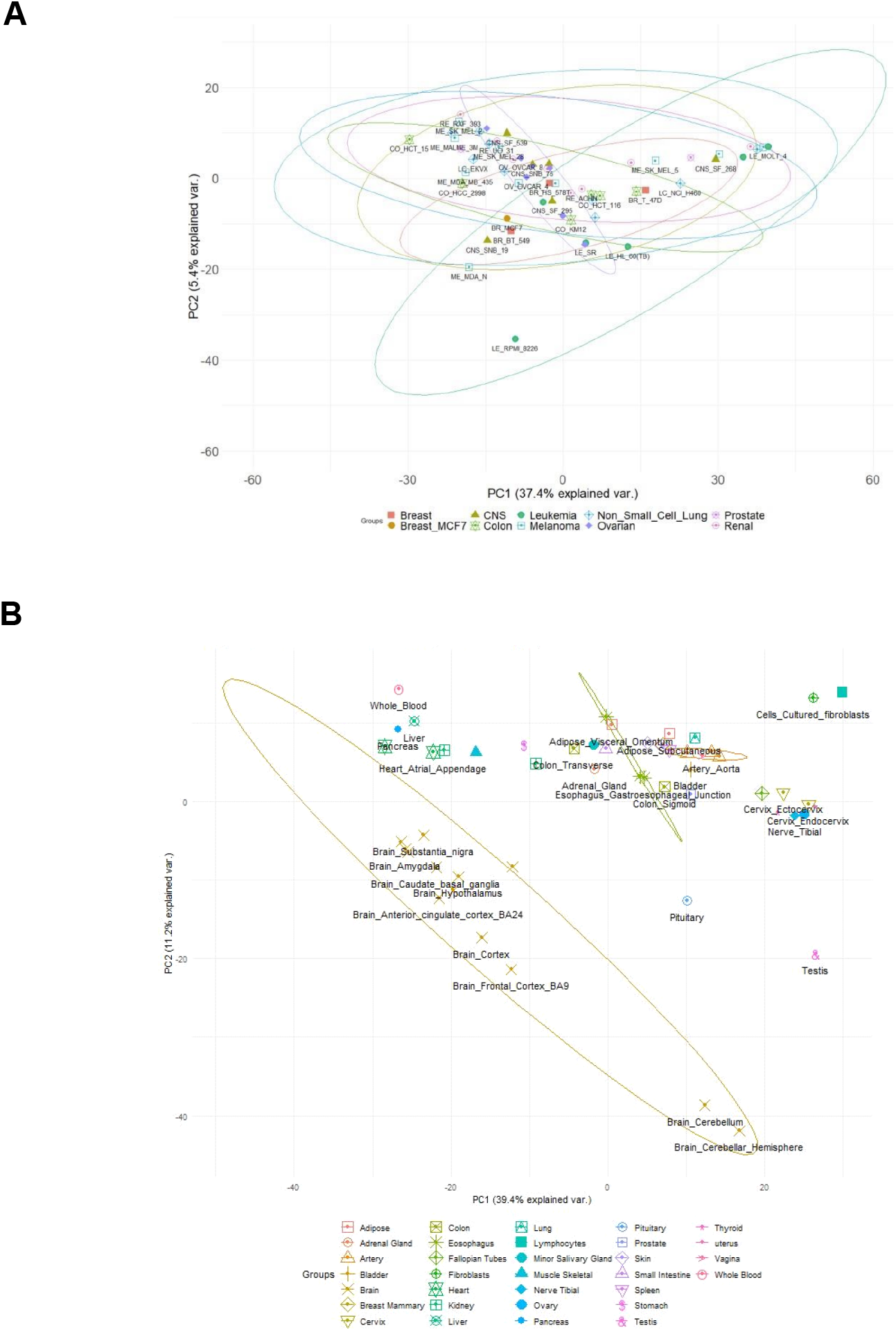
PCA plots of NCI-60 cancer cell lines (A) and GTEx normal human tissues (B) for the 9,753 genes expressed in all cell lines and all tissues. Two NCI-60 cell lines are excluded from (A), the breast cancer cell line MDA-MB-231 and the colon cancer cell line SW-620, as these two cell lines were widely separated in the PCA plot from their own congeners. Ellipses are 95% confidence ellipses for each group of cell lines or tissues. The 95% confidence ellipses for the cell lines all overlap and the majority of the cell lines are clustered in the center of the distribution of cell lines. In contrast, normal human tissues are displaced from each other. Even normal human brain tissues (14 tissues), while dispersed, are distinct from the other tissues. PCA analyses and plots generated with the statistical computer language R (ver. 4.1.0)

When comparing individual human tissues to cell lines, there are tissues that appear similar in their expression profile to several of the cancer cell lines. Figure 2A shows normal human liver tissue relative to the NCI-60 panel. Despite the fact the NCI-60 cell line panel does not include any human liver cancer cell lines, the GTEx normal human liver expression shows a profile like several of the cancer cell lines. A mix of melanoma, breast, non-small cell lung, and renal cancer cell lines appear closest to the GTEx human liver sample in the PCA plot. However, the overlapping 95% confidence ellipses for all the cancer cell lines indicate human liver expression is not very different from any of the cancer cell lines apart from ovarian cancer cell lines (the purple and smallest ellipse at the center of the distribution). This indicates there may be evidence that some cancer cell lines may behave more like unperturbed normal human liver tissue than others although all original tissue group confidence intervals overlap.

**Figure 2.**
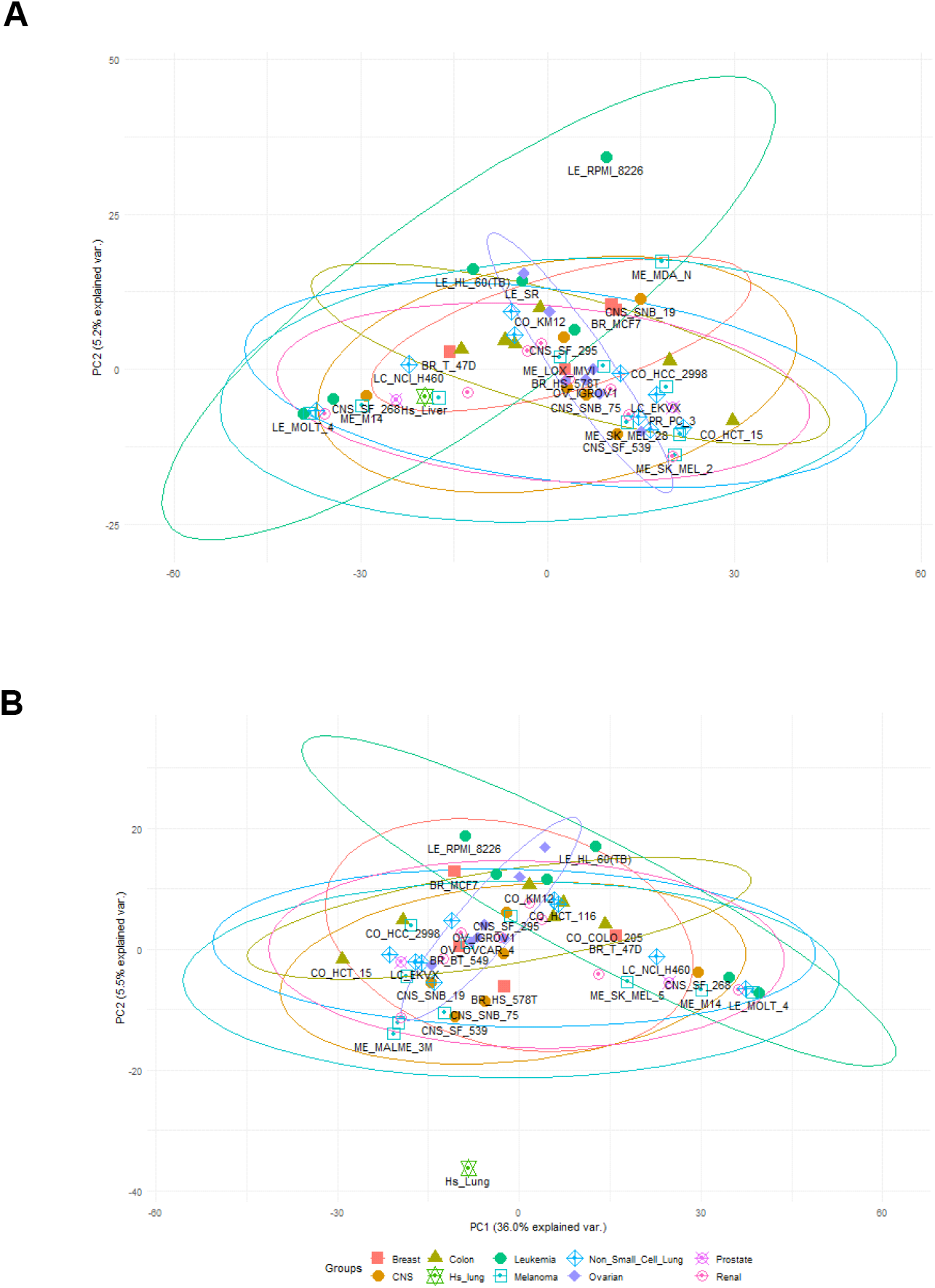
GTEx liver tissue relative to the NCI-60 cell lines for the 9,753 genes expressed in all human tissues and all NCI-60 cell lines. As in Figure 1, the breast cancer cell line MDA-MB-231 and the colon cancer cell line SW_620 have been excluded due to the apparent low expression of these two cell lines. (A) The NCI-60 cell line panel does not include any human liver cancer cell lines, yet normal human tissue expression appears not dissimilar from several cancer cell lines. (B) Conversely, a few GTEx human tissue expression profiles appear dissimilar to all NCI-60 cancer cell lines as shown in (B) for GTEx human lung tissue.

This differs from the GTEx normal human lung tissue (Figure 2B). The NCI-60 panel includes nine non-small cell lung cancer cell lines, and these are well scattered in the PCA plot (the light blue crossed diamond symbols). Normal human lung tissue is displaced from all the cancer cell lines. This pattern of normal human tissues being widely displaced from the cancer cell lines in the PCA analysis was also true for normal human colon, cervical, artery, and brain tissues. But other human tissues (kidney, spleen, skin, pancreas, prostate, stomach, testis, thyroid, uterus, vagina, whole blood, small intestine, tibial nerve, fallopian tube, breast, bladder, adipose tissue, colon, heart, muscle, and salivary gland) showed patterns similar to liver in which the GTEx expression profile was nested in the cancer cell line profiles but located much closer to some cell lines than others.

## DISCUSSION

Our intention in examining basal gene expression in cancer cell lines and normal human tissue was to determine sufficient differentiation in basal expression profiles to group cell lines as more or less likely to be reasonable surrogates for a given human tissue (or group of tissues) based on the pattern of expression in an unperturbed normal growing state. This would then aid in determining cell lines for a toxicity screening program and avoid unnecessary effort if fewer cell lines could be reasonably expected to cover the same biological space. This revelation is key to establishing a baseline of in vitro models for the use of NAMs in toxicity assessment of compounds.

Regarding basal gene expression profiles depending on original tissue, the NCI-60 cancer cell lines don’t seem to differ all that much. Furthermore, numerous cancer cell lines exhibit basal expression profiles that are very similar. The 95% confidence ellipses in Figures 1 and 2 indicate little difference between the cell lines in terms of their human tissue of origin. This would imply that in an unperturbed state, these immortalized cancer cell lines have very similar levels of expression. This was also true for the few genes expressed in all cancer cell lines but not expressed in the GTEx human tissue data. This small number of genes also showed a very tightly clustered PCA plot. Most of these genes appear to be related to cell cycle processes (when used as a gene list for ontology enrichment using Reactome, data not shown), so they presumably represent genes either essential to cancerous cell growth or related to processes of immortalization.

For decades, immortalized cancer cell lines have served as effective tools for in vitro efficacy testing and gross toxicity testing. However, it has long been known that they have several drawbacks or undesired characteristics for toxicity screening applications. Immortalized cell lines are convenient, but they are not truly biologically relevant (Roggen, 2011). Indeed, it is critical to have in vitro systems that adequately mimic key events of the in vivo mechanisms of action after exposure to a toxic compound. In fact, depending on the endpoint and type of chemicals to be evaluated, immortalized cancer cell lines may no longer display significant in vivo-like activity. According to the NCI-60 cancer cell lines’ basal gene expression, there isn’t much of a difference between cancer cell lines from very various origins as shown by the core set of genes that are expressed in all cell lines and all human tissues. This makes it challenging to assess a cancer cell line’s potential as a substitute for a certain type of human tissue. It is important to note that no cell lines generated from human liver cancer are included in the NCI-60 cancer cell line panel. However, a few of the cell lines in the NCI-60 panel have basal gene expression patterns that resemble very closely those found in the liver tissue of an average human. This may give some justification for choosing cell lines depending on how closely their normal expression resembles that of a tissue.

It appears that the basal expression profile of an immortalized cancer cell line is largely indicative of the fact that such cell lines, when simply growing in media, express genes to a similar degree. This is not a novel finding, as others have reported similar observations. Gillet et al. in studying the relevance of cancer cell lines to anti-cancer drug resistance observed that “all of the cell lines, grown either in vitro or in vivo, bear more resemblance to each other, regardless of the tissue of origin, than to the clinical samples they are supposed to model” (Gillet et al., 2011). Presumably this is an indicator that, while having different origins, there is a common underlying biology, derived from immortalization and carcinogenesis, that drives the dominant normal expression profile. It is then the way a cell line mounts a response to a chemical or physical perturbation that determines its specific suitability as a model for a given tissue response. To that end, toxicity screening programs may be limited in the range of relevant biological response captured using a panel of cancer cell lines. Cancer cell lines may be limited in response to perturbation due to the same limitations imposed on their transcriptome as a result of immortalization and being derived from cancerous tissues. In designing small scale screening projects for detailed MOA analyses or comparative or read-across analyses, it may be necessary to perform preliminary analyses on several cell lines to determine suitability for a specific application to distinguish if issues such as degree of metabolic competency fundamentally change assessment of MOA or POD values. To this end, comparison of cancer cell lines with primary cells or alternative in vitro models may be beneficial prior to a detailed in vitro dose response study for analyses of MOA.

Another approach is to turn to non-cancer cell lines already in widespread use. Human-induced pluripotent stem cell derived hepatocytes have been utilized for toxicology testing and drug screening (Davila et al., 2004; Mann, 2015). These have their own set of complications. One of the most notable is that the end cell population is inherently a mixed one, with cells expressing aspects of both adult and fetal transcriptomes. However, with transcriptomics, this is a knowable limitation as the genes from both developmental programs are known and can be identified from sequence data. Stem cell derived hepatocytes do display significant improvement in metabolic competence over cancer cell lines, and with refinement they may be a model replacement for primary cells in liver toxicity testing. Similarly, hTERT (human telomerase reverse transcriptase) immortalized cells have shown potential value in human toxicity testing for liver and bladder toxicity (Ramboer et al., 2015; Simon-Friedt et al., 2015; Wise et al., 2016; Smits et al., 2017).

Advances in next-generation sequence technology enable us to assess the transcriptomic behavior and suitability of an in vitro system as a surrogate for human tissue (Liu et al., 2019). While immortalized cancer cell lines have proven useful and remain so, it is becoming increasingly apparent that better model systems are needed and increasingly available. Transcriptomic analysis presented here is part of ongoing effort to end animal testing and develop a NAM framework using in vitro technologies for modern toxicology.

## METHODS

### Extraction of GTEx RNA-seq Datasets

The GTEx whole transcriptome RNA-Seq data is available for 54 human tissues sampled from more than 800 individuals (https://www.gtexportal.org/home/ and are available in the NCBI’s dbGaP database under accession phs000424.v8.p2). The expression data was provided as median transcripts-per million (TPM) expression values for basal (unperturbed) normal tissue (https://storage.googleapis.com/gtex_analysis_v8/rna_seq_data/GTEx_Analysis_2017-06-05_v8_RNASeQCv1.1.9_gene_median_tpm.gct.gz). TPM, as a measure of normalized expression, has value as a comparative measure of expression because it can be interpreted as “for every 1,000,000 RNA molecules in the RNA-seq sample, *X* of those came from gene *Y*.” This then provides a metric for direct comparison of expression across samples. A TPM value of zero in the data indicates a gene that was not detected at all, and thus assumed not to be expressed. This will inherently underestimate expression of genes expressed at very low levels (and thus with a very low probability of being detected), but in the absence of clear evidence for the detection of a gene, the assumption of no expression (or effectively, extremely low expression) is valid for these comparisons across tissues. The GTEx tissue list includes tissues relevant to toxicology target tissues such as liver, lung, multiple brain tissues, heart, thyroid, and kidney (https://gtexportal.org/home/datasets,) (Moore, 2013; Gibson, 2015).

### NCI-60 Cell Line Panel Basal Gene Expression

Whole transcriptome RNA-Seq data (dataset: ‘RNA:RNA-seq: composite expression’) from the National Cancer Institute’s NCI-60 Human Tumor Cell Line Screen are available through the CellMiner™ website (https://discover.nci.nih.gov/cellminer/loadDownload.do, see article PMID:31113817 for details to download data) (Shoemaker, 2006; Reinhold et al., 2019). This 60-cell line panel includes cell lines from nine cancer types (breast, CNS, colon, leukemia, melanoma, non-small cell lung, ovarian, prostate, and renal).

### Comparison of genes expressed in all GTEx human tissues and NCI-60 cancer cell lines as a basis for comparing normal constitutive expression in all cell lines and tissues

We determined the set of genes expressed in all human tissues and NCI-60 cell lines. A gene was considered as being actively expressed by detecting at least one RNA-Seq read per gene (for any transcript for that gene). These 9,753 genes represent those genes where a transcript was detected in all samples in the two datasets, and thus represent the most conservative gene set with which to compare the two sources of data. A principal component analysis (PCA) was performed, highlighting the distribution of the variation between the samples. We also generated 95% confidence ellipses for the cell lines all overlap and most of the cell lines are clustered in the center of the distribution of cell lines. Two of the cancer cell lines were excluded in this analysis as both were displaced from the other cell lines derived from the same human tissue. The breast cancer cell line MDA-MB-231 and the colon cancer cell line SW-620 had expression profiles that resulted in these two cell lines being displaced from the other cell lines and were removed to avoid any potential bias due to their unique expression signatures. The MDA-MB-231 cell line displays TPM values for many genes consistently lowest among all breast cancer cell lines or even lowest among all cancer cell lines. Similarly, the colon cancer cell line Sw-620 also shows a similar pattern of TPM values much different than the other six colon cancer cell lines and was similarly excluded from comparison.

## DATA AVAILABILITY

The processed and computed GTEx expression data is available for download in Mendeley Data with the access link: https://data.mendeley.com/datasets/9x7gyypvvk/1. The first tab of the table includes the downloaded transcripts per million (TPM) values of the NCI-60 cells; the third tab includes the downloaded median TPM values of the GTeX datasets, and the second tab contains the computed set of genes that overlap between the NCI-60 and GTeX datasets, and serves as the base gene list for the downstream analyses presented.

## CODE AVAILABILITY

The PCA plots in this study were generated by using the R software (v4.1.0) and the plotPCA package (https://rdrr.io/bioc/BiocGenerics/man/plotPCA.html).

## ACKNOWLEDGEMENTS

This work was supported by the American Chemistry Council’s Long-range Research Initiative.

## AUTHOR CONTRIBUTIONS

AN and MBB devised the project, MBB performed the analyses with input from PDM, ARB and AN, MBB, AN and ARB wrote the manuscript.

## COMPTETING INTERESTS

All authors are employed by ScitoVation LLC. All authors declare that the research was conducted in the absence of any commercial or financial relationships that could be construed as a potential conflict of interest.

